# Highly Accurate Estimation of the Fold Accuracy of Protein Structure Models

**DOI:** 10.64898/2026.04.15.718808

**Authors:** Lei Xie, Enjia Ye, Haodong Wang, Tianyou Zhang, Qihang Zhen, Fang Liang, Dong Liu, Guijun Zhang

## Abstract

**Background:** The function of a protein is intrinsically linked to its three-dimensional fold, and deep learning has revolutionized the field by enabling high-accuracy structure prediction at an unprecedented scale. Nevertheless, the growing deployment of these predictive pipelines in drug discovery and structural biology reveals a critical bottleneck that lies in the lack of independent and rigorous estimation of model accuracy (EMA) methodologies.

**Results:** Here we present DeepUMQA-Global, a single-model deep learning framework for estimating accuracy of protein structure models. Our method employs a structure-sequence cross-consistency mechanism to evaluate the bidirectional compatibility between the predicted structure and the input sequence, enabling comprehensive characterization of fold accuracy. DeepUMQA-Global outperforms the self-assessment confidence scores of AlphaFold3, achieving improvements of 57.8% in Pearson correlation and 49.0% in Spearman correlation. With respect to the CASP16 retrospective benchmark, DeepUMQA-Global outperforms all single-model accuracy estimation methods that participated in CASP16 and achieves performance comparable to that of the top consensusÖbased methods. A lightweight consensus strategy built upon DeepUMQA-Global ranks first among all CASP16 participants, surpassing all other methods, including consensus approaches, and highlighting the strength of our method. Remarkably, DeepUMQA-Global demonstrates a strong ability to discriminate between alternative conformational states of proteins, as evidenced in the CASP unique alternative conformation protein complex target and the CoDNaS benchmark.

**Conclusions:** Our results indicate that DeepUMQA-Global can be extended to broader protein modeling tasks, moving beyond static evaluation to offer a foundation for dynamic conformation EMA, where it accurately discriminates alternative conformational states and demonstrates reliable predictive fidelity in model accuracy estimation.

## Background

Estimation of model accuracy (EMA), also termed model quality assessment (MQA), is critical for the reliable interpretation and practical application of predicted protein structures in fields such as drug discovery, protein engineering, and functional analysis^1–3^. Without accurate EMA, errors originating from low-quality models can propagate into downstream experimental design and scientific conclusions, thereby undermining their validity^3,4^. Consequently, the development of highly accurate, robust, and generalizable EMA methods now represents both an urgent priority and a major focus in protein structure prediction, one that will ultimately determine the translational value and scientific impact of the current prediction revolution^5–7^.

Recent transformative advances, most notably AlphaFold2^8^ and AlphaFold3^9^, have spurred the development of a wide range of competing approaches for protein structure prediction, such as Boltz-2^10^, Chai-1^11^, SeedFold^12^, IntelliFold-2^13^, Protenix-v1^14^ and MassiveFold^15^, each capable of generating large ensembles of structural models. This unprecedented abundance has consequently shifted the field’s primary challenge from structure prediction to the reliable identification of the most accurate model among many candidates^16–18^. Accordingly, accurate EMA has emerged as a pivotal and pressing concern, marking a fundamental shift that reframes the core question from “Can we predict a structure?” to “How do we rank and select the best model?”^6,16–21^.

Model Accuracy Self-Assessment, which refers to the embedded confidence metrics generated by protein structure prediction methods to evaluate the reliability of their own predicted models, has been widely adopted by both deep learning-based predictors, including AlphaFold2^8^, AlphaFold-Multimer^22^, AlphaFold3^9^, and RoseTTAFold-All-Atom^23^, as well as physics-based docking methods such as HDOCK^24^. Generally, the predictive accuracy of protein monomer structures, particularly for single-domain proteins, is now approaching experimental resolution, underpinned by integrated self-assessment systems whose reliability matches experimental benchmarks^6,8,25^. This advancement delivers essential quality assurance for the practical deployment of predicted models across downstream applications^20^.

When the focus shifts to protein complex systems, the inherent challenges in the EMA of current methods become evident. Most existing self-assessment methods are derived from, or tightly integrated into, protein structure prediction frameworks that are strongly dependent on multiple sequence alignment (MSA) and are primarily designed for static protein structures^8,22^. This reliance is particularly pronounced in difficult scenarios, such as immune complexes with sparse co-evolutionary signals, large-scale macromolecular assemblies, and functionally relevant dynamic conformations^8,22,26,27^. Notably, the scores produced by these self-assessment methods are derived from the model’s own representational inference process, effectively reflecting internal confidence estimates of the generative model itself^8,24^. Consequently, their reliability and interpretability in crossÖmethod comparisons and in integrated validation frameworks remain substantially constrained. This limits their suitability as a generalizable and unbiased external standard for model quality assessment^19,28^.

These observations underscore the demand for dedicated, modelÖagnostic accuracy estimation methods. The growing recognition of this need is reflected in the latest Critical Assessment of Protein Structure Prediction (CASP) experiments (CASP15^29^ in 2022 and CASP16^30^ in 2024), which have introduced specific EMA evaluation tracks for protein complexes. These tracks benchmark and promote the most advanced third-party EMA methods, establishing them as the current state-of-the-art in the field. Based on the outcomes of the past two CASP rounds (CASP15 and CASP16), EMA methods are broadly classified into two principal categories: consensus-based and single-model approaches^16,17^. In parallel, a suite of additional derivative strategies has emerged, with quasi-single-model and meta-methods representing prominent examples^16,17^.

Consensus-based methods evaluate the accuracy of target models by integrating structural consistency among multiple candidate models. The core hypothesis is that regions exhibiting high structural consistency within a set of predicted models are generally more reliable. These methods typically perform “allÖagainstÖall” comparisons across an input model pool and generate quality scores based on statistical or geometric consistency metrics (e.g., pairwise similarity scores or structural clustering)^16–18,29,31^. Representative approaches include Pcons^32^, VoroIFÖjury^33^, the ModFOLDdock series^34,35^, DeepUMQAÖX^36^, and the MULTICOM series^37,38^. While consensus-based approaches can deliver highly accurate estimates, their effectiveness is critically dependent on the quality and diversity of the input model pool. When the pool lacks sufficient high-quality predictions, the reliability of assessment declines markedly^6,7,29,39^. In cases where models are highly similar, consensus methods struggle to discriminate subtle quality differences. Furthermore, for flexible regions or proteins exhibiting multiple conformational states, consensus-based approaches may tend to underestimate accuracy, as conformational diversity is often misinterpreted as structural inconsistency^40,41^. It is worth noting that some approaches, although classified as “quasiÖsingleÖmodel” methods (e.g., MIEnsemble^31^ and ModFOLDdock2S^35^), internally generate diverse models and perform consensusÖlike comparisons, thereby retaining strong consensus characteristics. This illustrates the extensibility and integrability of consensus strategies across different methodological frameworks.

The fundamental principle of single-model methods is to conduct accuracy estimation based exclusively on the features of an individual predicted model, without reliance on external model sets or consensus comparisons. Representative approaches in this area include the ProQ series^42–44^, DeepAccNet^45^, VoroIF-GNN^46^, and the DeepUMQA series^47–49^. It was noted in the CASP16 assessors’ summary report^16^, that the highest-performing true single-model method was our team’s GuijunLab-PAthreader, which is built upon the previously developed DeepUMQA3^49^ protocol. Single-model methods demonstrate distinct advantages, including independence for use in single-model scenarios, high computational efficiency without requiring multi-model alignment, and strong interpretability derived from direct analysis of the model’s structural or statistical features. Even when the overall quality of the model set is constrained, single-model methods remain robust for assessing performance^16,17^. Meanwhile, meta-methods have continued to evolve, advancing beyond simple geometric consensus toward the strategic integration of results from multiple single-model approaches using machine learning techniques^16^. This strategy allows for enhanced overall quality assessment, as demonstrated by methods such as ModFOLDdock2^35^. Such progress further highlights the indispensable role of single-model methods as core assessment units^21^. They not only provide the foundational input for meta-method integration, but also represent a vital pathway for achieving efficient and reliable evaluation under resource-constrained scenarios.

Single-model EMA methods are widely regarded as the “holy grail” of the field, as they embody the original and fundamental goal of EMA: estimating model accuracy based on a single predicted structure^17^. In recent years, these methods have achieved significant progress, particularly in the evaluation of local residue accuracy^17^. However, when tasked with assessing the global assembly quality of protein complexes, single-model methods struggle to outperform consensus strategies that rely on multi-model comparisons (e.g., fold and interface scores). Meanwhile, single-model methods have yet to surpass the accuracy level of the built-in self-assessment metrics of the AlphaFold series^21^. These defects highlight a performance bottleneck in single-model EMA approaches. Consequently, bridging this gap to match or even surpass the performance of self-assessment or consensus-based methods remains a pivotal challenge^21^. Leveraging the deviation of the bidirectional consistency between sequence and structure may offer a promising pathway to cope with this challenge.

In this work, we introduce DeepUMQA-Global, a single-model method designed for evaluating global accuracy. Building upon our previously developed DeepUMQA3 algorithm^49^, this approach exploits the bidirectional compatibility between a target’s sequence and its three-dimensional structure, enabling accurate evaluation of a model’s global quality through a structureÖsequence cross-consistency mechanism. Experimental results show that DeepUMQA-Global substantially outperforms AlphaFold3’s self-assessment, achieving 57.8% and 49.4% higher Pearson and Spearman correlation coefficients, respectively. Meanwhile, DeepUMQA-Global substantially outperformed all the other single-model methods in the CASP16 benchmark test set and exhibited performance comparable to that of the consensus-based approaches. Importantly, DeepUMQA-Global demonstrates both the ability and the potential to reliably assess and discriminate alternative conformational states.

## Results and Discussion

### DeepUMQA-Global Overview

DeepUMQA-Global is a deep learning framework for estimating the fold accuracy of protein structural models at both the global complex (pScore) and interface (ipScore) levels (**Fig. 1**). We introduce a structure–sequence cross-consistency mechanism that captures the bidirectional compatibility between the three-dimensional structure and the amino acid sequence.

**Fig. 1.**
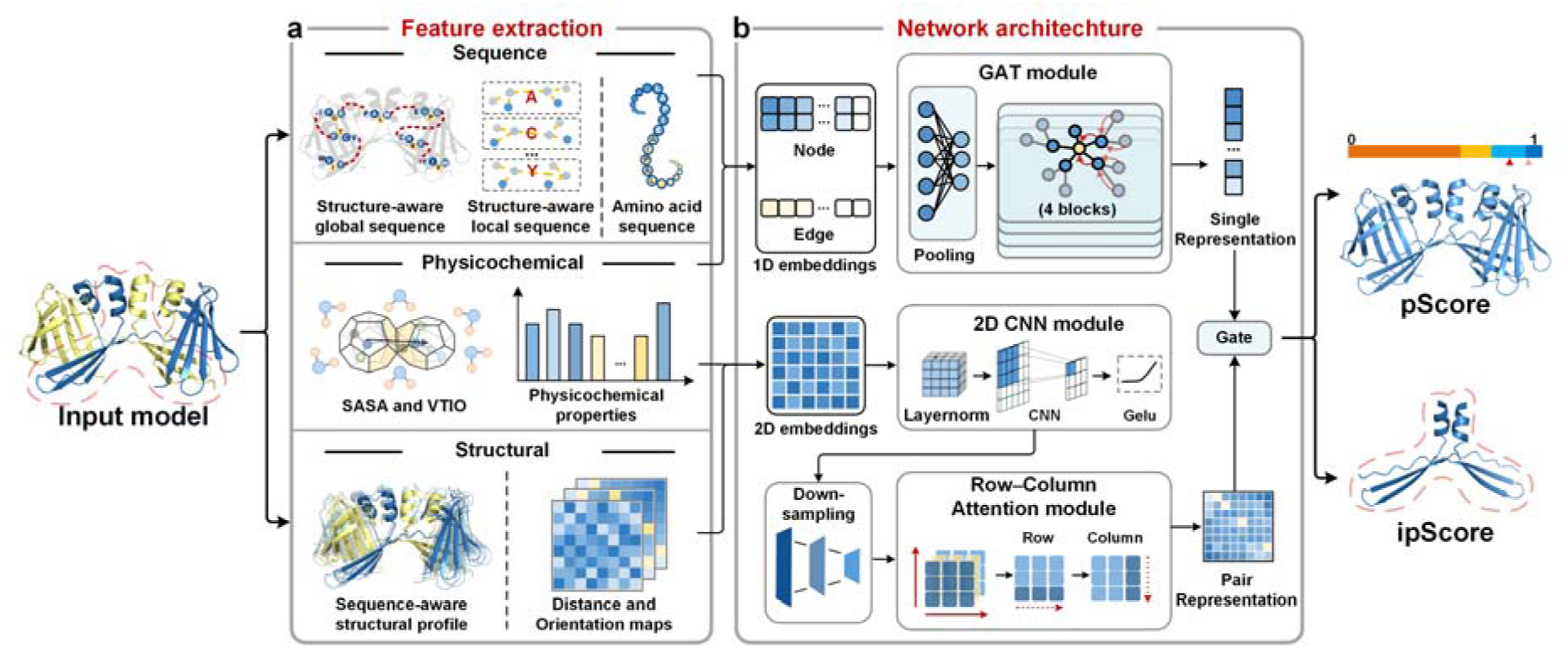
Overview of DeepUMQA-Global. DeepUMQA-Global evaluates an input protein model’s accuracy by predicting the complex global score (pScore) and interface global score (ipScore). **a,** Feature extraction. DeepUMQA-Global extracts three complementary feature categories from each input model, including sequence features (structure-aware global sequence (SAGS), structure-aware local sequence (SALS), and input amino acid sequence), structural features (sequence-aware structural profiles together with residue–residue distance and orientation maps), and physicochemical features (solvent-accessible surface area (SASA), Voronoi-tessellation interface orientation (VTIO), and physicochemical properties that include meiler, amino acid properties, BLOSUM62, Rosetta one-body and two-body energy terms, backbone bond lengths and angles, torsion angles, Euler angles and voxelization). **b,** Residue-wise features are concatenated as nodes, with edges connecting residue pairs whose C atoms are within 15 Å, collectively forming 1D embeddings that are encoded by a graph attention network (GAT) module to produce the single representation. In parallel, the inter-residue features are concatenated into 2D embeddings, and processed by the 2D convolutional neural network (2D CNN) and Row–Column attention module to generate the pair representation. The single representation gates the pair representation to produce the final complex and interface global score.

In the feature extraction stage (**Fig. 1a**), DeepUMQA-Global extracts three complementary types of features from each the input model, including sequence, structure and physicochemical features. Sequence features comprise the amino acid sequence of input model together with two structure-informed sequence representations, referred to as structure-aware global sequence (SAGS) and structure-aware local sequence (SALS). (1) SAGS is derived from embedding representations generated by ProteinMPNN^50^, which predicts sequence distributions conditioned on the three-dimensional structure, encoding global structure-conditioned sequence preferences that are subsequently provided to the network for learning structure–sequence consistency patterns. SALS is obtained from 3Di descriptors generated by Foldseek-Multimer^51,52^, which encode local structural environments as discrete sequence-like tokens, providing structure-conditioned representations of residue contexts. (2) Structural features comprise a sequence-aware structural profile (SASP) together with residue–residue distance and orientation maps. SASP encodes template-derived residue–residue distance profiles, in which inter-residue distances are represented as distributions accumulated across structurally aligned templates. These templates are retrieved using the AlphaFold3-predicted structure generated from the sequence of input model as the search query. The accompanying entropy term captures residue–residue uncertainty across templates, and SASP provides structural priors. In contrast, the distance and orientation maps are directly computed from the input structure. Together, these features enable the network to assess structural consistency by comparing the input model against sequence-conditioned topological constraints. Notably, SAGS and SASP together provide a bidirectional view of sequence–structure compatibility: the former reflects structure-conditioned sequence preferences, whereas the latter captures sequence-conditioned structural priors. (3) Physicochemical features consist of solvent-accessible surface area (SASA) and voronoi tessellation-derived interface orientation (VTIO)^46,53^, together with physicochemical properties, including meiler^45^, amino acid properties, BLOSUM62^54^, Rosetta one-body and two-body energy terms^55^, backbone bond lengths and angles, torsion angles, Euler angles and voxelization. SASA encodes per-residue surface exposure by quantifying both the total contact area with neighboring residues and the solvent-exposed area. VTIO represents a Voronoi tessellation-derived contact orientation vector that captures the spatial bias of inter-residue contacts, thereby describing local packing directionality. Physicochemical properties further characterize the physical and chemical plausibility of the model. Detailed shapes and descriptions of all features are provided in **Supplementary Table 1**, with the specific feature extraction and encoding procedures described in the **Method Details** section.

The network architecture (**Fig. 1b**) processes residue-wise and inter-residue features in parallel. Residue-wise features are concatenated as node features, and edges are defined for residue pairs whose C_α_ atoms are within 15OÅ, together forming one-dimensional embeddings. These embeddings are processed by a graph attention network (GAT)^56^ module to generate the single representation, which integrates amino acid sequence information with structure-aware sequence features to capture residue-level sequence–structure consistency. In parallel, inter-residue features are concatenated into two-dimensional (2D) embeddings, which are processed by the 2D convolutional neural network (2D CNN)^57^ module and the Row–Column attention^58^ module to generate pair representation, which combines geometric features derived from the input structure with sequence-aware structural priors, enabling the model to assess inter-residue topological consistency. The single representation is broadcast and applied via element-wise multiplication to gate the pair representation, enabling the network to preferentially aggregate inter-residue signals supported by locally reliable regions. The gated representations are averaged over both rows and columns, effectively integrating residue–residue interactions across the entire structure to generate the complex global accuracy score (pScore) and the interface global score (ipScore). For the ipScore, only residues at the protein–protein interface (defined as those with at least one inter-chain atom within 5 Å) are considered, and the same network pipeline is applied to these residues. The network is trained under the supervision of the TM-score^59^ to capture overall model accuracy. Details of the network architecture and training procedure are provided in the **Methods Details**.

Single-model methods are inherently limited by the information available in a single predicted structure. Unlike self-assessment approaches, they cannot access representations generated during the structure prediction process, and unlike consensus methods, they do not leverage consensus information across multiple candidate models. To address this limitation, DeepUMQA-Global couples a bidirectional sequence–structure compatibility signal within a single model, integrating structure-aware sequence plausibility and sequence-aware structure consistency to enable robust global accuracy evaluation within a single-model framework. The test set constructed from AlphaFold3-generated models, results indicate that, as an independent single-model EMA method, DeepUMQA-Global substantially outperforms AlphaFold3’s self-assessment in evaluating both complex and interface global accuracy, while achieving performance on the CASP16 benchmark test set that is comparable to consensus-based approaches. Moreover, the performance on proteins with alternative conformational states highlights its potential to evaluate and discriminate distinct conformations.

### Performance Comparison of DeepUMQA-Global with AlphaFold3 Self-Assessment and Docking-based Self-Assessment Methods

To facilitate direct performance comparison with the AlphaFold3 self-assessment, we constructed a test set based on 38 targets from CASP16. Given the computational limitations of AlphaFold3 in handling large assemblies, only targets with fewer than 3,000 residues were retained. Additionally, two dynamic targets (T1249 and T1294) were excluded to minimize potential confounding effects arising from conformational heterogeneity. This filtering process yielded a final test set of 26 protein complex targets. The training set consisted of 7,840 monomeric Protein Data Bank (PDB)^60^ targets collected before March 1, 2022, each generating more over 100 decoys, resulting in a total of more than one million models^45^. To ensure a strict separation between the training and test sets, two criteria were applied as follows. First, all test targets were selected from CASP16, whose native structures were released in May 2024. Second, any training target with sequence identity greater than 40% to the selected test targets was removed, guaranteeing that the test proteins were non-redundant with respect to the training set. Since AlphaFold3 confidence scores are only available for structures generated within its own inference pipeline, we generated corresponding decoys for each target using the AlphaFold3 online server (https://alphafoldserver.com/). For each of the 26 targets, decoys were generated stochastically using random seeds, producing 50 decoys per target. For comparison with docking-based self-assessment, a test set was constructed from dimeric PDB targets deposited between January 1 and April 1, 2024. Targets with resolution better than 2.5OÅ and chain lengths of 20–1,000 residues were included. After 40% sequence redundancy removal within the set and against the training set, 107 dimeric complexes were retained. For each dimeric complex, the component monomers were re-docked using HDOCK^61^ and DiffDock-PP^62^. For each method, 200 decoys were generated, producing two separate sets of decoys per target. The detailed procedures for dataset construction are provided in the **Method Details** section.

In this study, we used the TM-score^59^, which was calculated between the predicted models and their corresponding experimentally determined reference structures, as the reference metric for evaluating structural model accuracy. To comprehensively evaluate the estimation performance of each method, we used three statistical metrics: Pearson and Spearman correlation coefficients, which quantify the linear and monotonic agreement, respectively, between the predicted scores and the ground-truth TM-scores, as well as the ROC AUC, which assesses the capability of each method to discriminate high from low-quality models, where high-quality models were defined as the top 25% of all models.

We first benchmark the performance of DeepUMQA-Global against the AlphaFold3 self-assessment on our constructed test set. DeepUMQA-Global shows substantially improved agreement with the reference structural accuracy compared with AlphaFold3, achieving the Pearson correlation coefficient of 0.453 (95% confidence interval = 0.340–0.566), the Spearman correlation coefficient of 0.371 (95% confidence interval = 0.269–0.473), and the ROC AUC of 0.614 (95% confidence interval = 0.558–0.670), compared with 0.287 (95% confidence interval = 0.200–0.375), 0.249 (95% confidence interval = 0.158–0.340), and 0.568 (95% confidence interval = 0.525–0.611) for AlphaFold3, respectively. These improvements correspond to gains of 57.8%, 49.0% and 8.1% in the Pearson and Spearman correlation coefficients and ROC AUC, respectively. The Pearson and Spearman correlation coefficients show statistically significant improvements, whereas the ROC AUC exhibits a smaller yet consistent gain (**Fig. 2a**), as confirmed by a two-sided Wilcoxon signed-rank test (P = 5.01×O10⁻^3^, 1.76×O10⁻^2^, and 1.33×O10⁻^1^, respectively). Detailed results are provided in **Supplementary Table 2**. Per-target analysis further supported the robustness of DeepUMQA-Global, which achieved better Pearson and Spearman correlation coefficients than AlphaFold3 for 18 out of 26 targets (69.2%) (**Fig. 2b, c**). Notably, we also observed that DeepUMQA-Global provides modest yet consistent improvements over AlphaFold3 on predicted protein complex interfacial regions (defined as inter-chain atomic contacts within 5 Å, according to the parameter settings used by AlphaFold3). On these regions, DeepUMQA-Global achieved higher Pearson and Spearman correlation coefficients, with gains of 38.8% and 22.9%, respectively, and a marginally improved ROC AUC, although these differences did not meet the threshold for statistical significance (**Fig. 2d**). Detailed results are provided in **Supplementary Table 3**. Per-target analysis further revealed that, among the 26 targets, DeepUMQA-Global outperforms AlphaFold3 in the Pearson correlation coefficient for 16 targets (61.5%), while showing comparable performance in the Spearman correlation coefficient for 13 targets (50%) (**Fig. 2e, f**). We consider that the improved performance of DeepUMQA-Global may be attributable to differences in training strategies compared with AlphaFold3’s internal confidence head. Specifically, DeepUMQA-Global is trained directly on large and diverse decoy sets that span a broad range of structural qualities, with optimization explicitly targeted toward correlation with global accuracy and reliable model ranking. In contrast, AlphaFold3’s confidence head is trained on models generated from limited mini-rollout^9^ during training, which may not fully represent the diverse-quality structure models encountered in practical evaluation. These results underscore the potential of DeepUMQA-Global, as an independent model accuracy estimation method, to serve as a useful complement to the built-in confidence scores of existing protein structure prediction approaches.

**Fig. 2.**
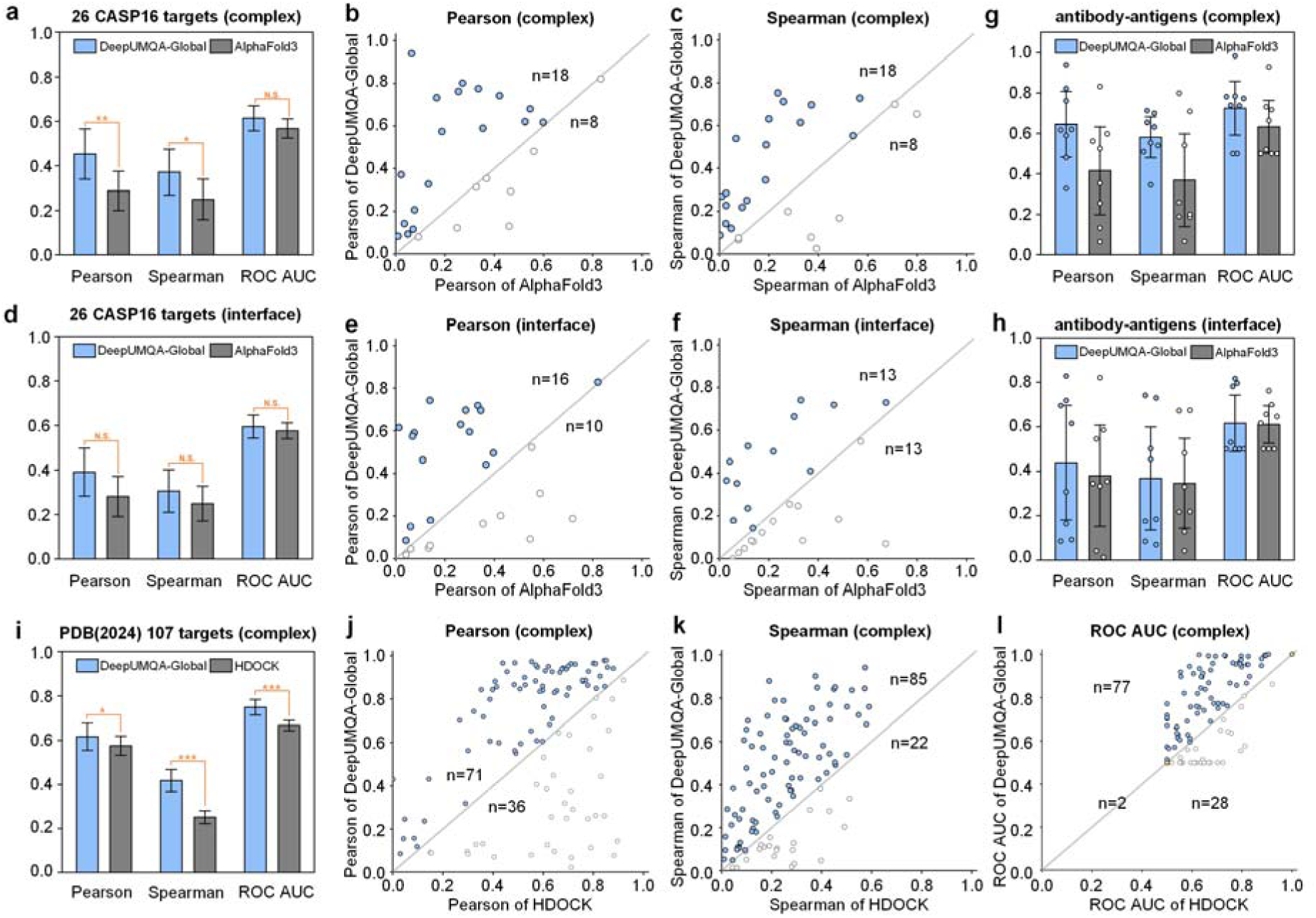
Performance comparison of DeepUMQA-Global with AlphaFold3 self-assessment and Docking-based self-assessment methods. a, Performance comparison between DeepUMQA-Global and AlphaFold3 self-assessment methods on 26 CASP16 targets. The bar height indicates the mean. The error bars represent 95% confidence intervals based on 10,000 bootstrap resamples across targets. Significance was assessed using the two-sided Wilcoxon signed-rank test on paired per-target metrics across the 26 targets for Pearson, Spearman correlation coefficients and ROC AUC (*P* = 5.01× J10⁻^3^, 1.76× 10⁻^2^, and 1.33× 10⁻^1^); \*\**P*<0.01, \**P*<0.05, and *N.S.* indicates no significance. **b,** Per-target comparison of Pearson correlation coefficients between DeepUMQA-Global and AlphaFold3 self-assessment, with each point representing one target. **c,** Per-target comparison of Spearman correlation coefficients between DeepUMQA-Global and AlphaFold3 self-assessment. **d,** Performance comparison between DeepUMQA-Global and AlphaFold3 self-assessment for the interface regions on the 26 CASP16 targets. **e,** Per-target comparison of the Pearson correlation coefficient between DeepUMQA-Global and AlphaFold3 self-assessment for the interface regions. **f,** Per-target comparison of the Spearman correlation between DeepUMQA-Global and AlphaFold3 self-assessment for the interface regions. **g,** Performance comparison between DeepUMQA-Global and AlphaFold3 self-assessment on eight antibody-antigen targets. **h,** Performance comparison between DeepUMQA-Global and AlphaFold3 self-assessment on the interface regions of eight antibody–antigen targets. **i,** Performance comparison between DeepUMQA-Global and HDOCK confidence scores on PDB(2024) 107 targets. Significance was assessed using a two-sided Wilcoxon signed-rank tests for the Pearson, Spearman correlation coefficients and ROC AUC (*P* = 3.87×10⁻^2^, 8.43×10⁻^11^ and 3.18× 10⁻^7^, respectively); \*\*\**P*<0.001, \**P*<0.05. **j,** Per-target comparison of the Pearson correlation coefficient between DeepUMQA-Global and HDOCK. **k,** Per-target comparison of the Spearman correlation between DeepUMQA-Global and HDOCK. **l,** Per-target comparison of the ROC AUC between DeepUMQA-Global and HDOCK.

Among the 26 targets in our constructed test set, eight are antibody–antigen complexes, which pose notable challenges due to the inherent flexibility of antibody complementarity-determining regions (CDRs)^17^. Given the distinct complexity and biological significance of such targets, we performed a focused analysis comparing DeepUMQA-Global and AlphaFold3 self-assessment on these eight CASP16 antibody-antigen targets. DeepUMQA-Global achieves a Pearson correlation coefficient of 0.645 (95% confidence interval = 0.483–0.807), a Spearman correlation coefficient of 0.581 (95% confidence interval = 0.480–0.682) and an ROC AUC of 0.725 (95% confidence interval = 0.592–0.858), compared with 0.415 (95% confidence interval = 0.198–0.632), 0.369 (95% confidence interval = 0.139–0.599) and 0.633 (95% confidence interval = 0.502–0.764) for AlphaFold3, respectively, with corresponding improvements of 55.4%, 57.5% and 14.5% (**Fig. 2g**). Given the limited size of this subset, we do not report significance test statistics here, and detailed results are provided in **Supplementary Table 4**. We also observed that DeepUMQA-Global exhibits modest performance gains over AlphaFold3 for the accuracy estimation of antibody–antigen interfacial regions (**Fig. 2h**), with detailed results provided in **Supplementary Table 5**. This modest performance gain at antibody–antigen interfaces is in marked contrast to the more substantial improvements seen for general protein complexes. The difference can be attributed to the unique biophysical properties of immune recognition, including weak coevolutionary signals, high conformational flexibility of the CDR loops, and a relatively flat binding energy landscape, all of which hinder accurate evaluation of interfacial regions. These results also suggest a need for more advanced evaluation frameworks for antibody–antigen complexes, moving beyond static structure-based approaches to incorporate dynamic conformational analysis or complementary biophysical descriptors, such as solvation free energy and side-chain entropy.

To further evaluate the robustness and general applicability of DeepUMQA-Global, we performed a dedicated benchmarking experiment against representative docking-based self-assessment approaches. To this end, we first compared its performance with the confidence score from HDOCK^61^, a widely used hybrid protein–protein docking method. The HDOCK confidence score, derived from knowledge-based docking potentials, serves as a baseline to assess the reliability of docked complex models. DeepUMQA-Global achieves a mean Pearson correlation coefficient of 0.616 (95% confidence interval = 0.555–0.677), a Spearman correlation coefficient of 0.418 (95% confidence interval: 0.368–0.468), and an ROC AUC of 0.749 (95% confidence interval: 0.715–0.783), compared with 0.575 (95% confidence interval = 0.531–0.619), 0.251 (95% confidence interval = 0.222–0.280), and 0.667 (95% confidence interval = 0.644–0.690) for HDOCK, respectively. These improvements are statistically significant (**Fig. 2i**), corresponding to gains of 7.1%, 66.5%, and 12.3%, for which significance was assessed using a two-sided

Wilcoxon signed-rank tests for Pearson, Spearman and ROC AUC (*P* = 3.87×10⁻^2^, 8.43×10⁻^11^ and 3.18× 10⁻^7^, respectively). Detailed results are provided in **Supplementary Table 6**. Per-target analysis revealed that DeepUMQA-Global outperforms HDOCK on 71 targets (66.4%) for Pearson correlation coefficients, 85 targets (79.4%) for Spearman correlation coefficients, and 77 targets (72.0%) for ROC AUC out of all 107 targets (**Fig. 2j**–**l**). We further compared DeepUMQA-Global with DiffDock-PP^62^, a deep learning-based docking method that provides confidence scores based on C_α_ - RMSD ^63^ (**Supplementary Fig. 1**). DeepUMQA-Global’s top-ranked models yield higher TM-score and lower C_α_ - RMSD relative to the native structure, demonstrating more accurate selection of near-native conformations. Detailed results are summarized in **Supplementary Tables 7 and 8**. Notably, the confidence scores of HDOCK and DiffDock-PP rely on the sampling of candidate models during their respective docking processes within individual targets. In contrast, DeepUMQA-Global serves as a unified evaluator capable of cross-method and cross-target model accuracy estimation. This distinctive feature eliminates the need for method-specific sampling and enables a fair, direct comparison of model quality across different docking pipelines and targets.

### Performance Comparison of DeepUMQA-Global with the CASP16 EMA Methods

To rigorously and consistently evaluate the performance of DeepUMQA-Global for global accuracy estimation, we perform a retrospective benchmark study following the evaluation protocol of the CASP16 EMA experiment^16^. This experiment provides blind test targets and standardized metrics specifically designed for assessing EMA methods. The CASP16 benchmark test set is composed of the models submitted by participating structure prediction groups for the official CASP16 target proteins. Global accuracy estimation was evaluated through two dimensions of the CASP16 EMA QMODE1 framework, SCORE and QSCORE, to assess the overall structural topology and the accuracy of the inter-chain interface of predicted complexes, respectively^16^. The official CASP16 EMA assessment included 38 targets for the SCORE evaluation and 39 for the QSCORE evaluation^16^. Two targets were excluded from the official assessment because their reference structures were not released. We further restricted the analysis to targets with sequence lengths less than 3,000 residues due to computational constraints, resulting in a final set of 30 target models for the CASP16 benchmark test set. According to the CASP EMA assessment criteria^16^, each method is required to provide predicted accuracy scores for at least 80% of the targets and to cover at least 80% of the submitted models within each target, and DeepUMQA-Global satisfies these coverage requirements. Analyses were conducted using the official CASP16 EMA pipeline (https://git.scicore.unibas.ch/schwede/casp16_ema).

For the SCORE evaluation, two global structural similarity metrics based on structure superposition were employed, including the TM-score^59^, which measures length-normalized global structural similarity between predicted and reference models, and Oligo-GDTTS^17^, which evaluates the global structural agreement of multimeric assemblies under rigid-body superposition. For the QSCORE evaluation, interface-specific metrics were used, including the QS-score^17^, which quantifies the agreement of interfacial residue contacts and interface topology between predicted and reference complexes, and DockQ-wave^17^, a continuous variant of the DockQ^64^ metric, which integrates the interface RMSD, ligand RMSD and fraction of native contacts into a single docking quality score. Four statistical metrics were used to comprehensively evaluate the consistency between the predicted accuracy scores and experimental reference values for both SCORE and QSCORE, including the Pearson correlation coefficient (P) for linear correlation, the Spearman correlation coefficient (S) for rank correlation, the ROC AUC (R) for discriminating high-quality models (defined as the top 25% best models) and low-quality models, and Loss (L) for quantifying the difference between the reference score of the model ranked highest by the estimator and the best reference score observed among all the candidate models^16^. Lower loss indicates better performance, whereas after normalization, higher Loss corresponds to superior performance. All four per-target statistical metrics were converted to Z-scores, with each metric standardized across all participating predictors on a per-target basis using the mean and standard deviation of the corresponding metric distribution, and negative Z-scores set to 0^16^. These Z-scores for the four statistical metrics were then combined for each reference metric following the predefined ranking scheme to generate a composite score per reference metric for each estimator, and final rankings for the SCORE and QSCORE evaluation were obtained by aggregating the corresponding composite Z-scores across their respective reference metrics^16^. Detailed definitions of the reference metrics and Z-score calculations are provided in **Supplementary Note 1**.

As DeepUMQA-Global is a single-model method, we benchmarked its performance against other CASP16 single-model EMA methods in a retrospective evaluation. Specifically, DeepUMQA-Global achieved the highest overall Z-scores of 109.568 in the SCORE evaluation and 108.718 in the QSCORE evaluation (**Fig. 3a**), exceeding the second-ranked single-model method by 38.285 and 11.180 Z-score units, respectively (**Supplementary Tables 9 and 10**). DeepUMQA-Global exhibits strong performance in Spearman, ROC AUC, and Loss, reflecting its strong ability to rank models, distinguish high- and low-quality models, and select near-native models from candidate models. In addition, DeepUMQA-Global achieved a Top1Loss of <0.05 in the SCORE evaluation for 16 targets (**Fig. 3b**), indicating its strong ability for top-model selection. Nevertheless, relatively weak Pearson correlation coefficients are observed, partly because of a slight overestimation of model accuracy, indicating that the room for improvement in quantitative absolute quality estimation remains. These results demonstrate that DeepUMQA-Global produces robust model accuracy estimation across diverse targets.

**Fig. 3.**
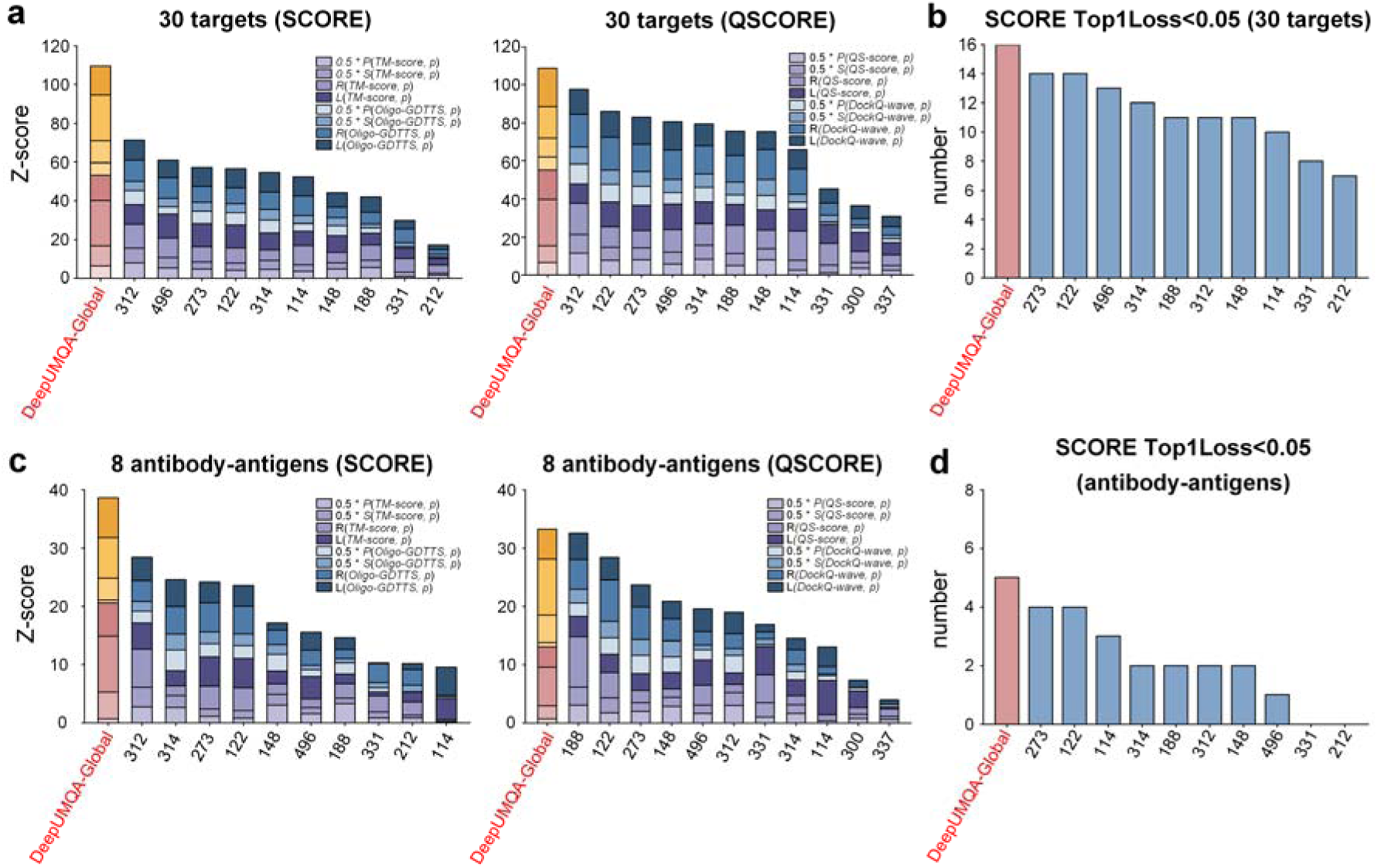
Performance comparison of DeepUMQA-Global with CASP16 single-model EMA methods. a, Performance comparison of DeepUMQA-Global with the CASP16 single-model EMA methods in terms of overall Z-score in the SCORE (left) and QSCORE (right) evaluations across 30 CASP16 targets. Overall Z-scores were computed by first calculating per-target Z-scores for each statistical metric (Pearson correlation, Spearman correlation, ROC AUC, and Loss), and then aggregating them for SCORE (based on TM-score and oligo-GDTTS) and QSCORE (based on QS-score and DockQ-wave) following the CASP16 EMA evaluation framework. **b,** Number of CASP16 targets with a Top1Loss of <0.05 in the SCORE evaluation, comparing DeepUMQA-Global with the CASP16 single-model EMA methods. **c,** Performance comparison of DeepUMQA-Global with the CASP16 single-model EMA methods in terms of overall Z-scores in the SCORE (left) and QSCORE (right) evaluations across eight antibody-antigen targets. **d,** Number of antibody-antigen targets with a Top1Loss of <0.05 in the SCORE evaluation.

To assess the performance on antibody-antigen targets, we evaluated DeepUMQA-Global on this challenging subset, characterized by conformational variability and flexible CDR loops (**Fig. 3c**). DeepUMQA-Global achieved the highest overall Z-score of 38.590 in the SCORE evaluation among all single-model EMA methods, outperforming the second-ranked method by 10.193 Z-score units (**Supplementary Table 11**). A similar performance trend was observed for QSCORE, where DeepUMQA-Global also achieved the highest overall Z-score of 33.315 (**Supplementary Table 12**). In addition, DeepUMQA-Global satisfied the SCORE evaluation top-model selection criterion (Top1Loss < 0.05) for five targets, surpassing all other single-model methods (**Fig. 3d**). These results indicate that DeepUMQA-Global retains robust performance on antibody-antigen complexes, enabling relatively reliable model ranking and accurate identification of near-native models despite pronounced interface flexibility.

The improved performance of DeepUMQA-Global in the CASP16 EMA QMODE1 can be attributed to its bidirectional sequence–structure cross-consistency framework, enabled by the synergistic integration of two key complementary features, sequence-aware structural profile (SASP) and structure-aware global sequence (SAGS). SASP introduces sequence-conditioned structural priors that constrain global fold organization and long-range residue–residue geometry, and SAGS captures the compatibility between the structure-aware sequence and the amino acid sequence of the input model. Together, these components impose complementary constraints on sequence–structure consistency, effectively reducing structurally implausible conformations and sequence–structure mismatches. This design may provide a plausible explanation for the robust model ranking and top-model selection performance observed for the CASP16 blind targets. Ablation analyses in the subsequent sections further support the individual and synergistic contributions of SASP and SAGS to the observed performance improvements.

Consensus EMA methods typically outperform single-model methods in CASP evaluations. The best-performing single-model method in CASP16 ranked only 10th in SCORE and 9th in QSCORE, indicating a substantial performance gap^16^. Despite this, single-model methods remain important due to their independence from multiple predictions and broader applicability in practical scenarios. To evaluate the competitiveness of DeepUMQA-Global under this background, we benchmarked DeepUMQA-Global against all EMA methods reported in CASP16, including consensus, quasi-single-model and single-model methods (**Supplementary Fig. 2**). DeepUMQA-Global achieved the second-highest overall Z-score in the SCORE evaluation (**Supplementary Table 13**) and the seventh overall Z-score in the QSCORE evaluation (**Supplementary Table 14**). Notably, DeepUMQA-Global achieved the highest Loss Z-scores for the TM-score, Oligo-GDTTS, and DockQ-wave, with strong Loss performance also observed for the QS-score, indicating its robust ability to select the top-ranked model. Furthermore, compared with other methods, DeepUMQA-Global achieved the highest Spearman and ROC AUC with respect to Oligo-GDTTS, indicating strong performance in rank ordering and model discrimination at the oligomeric assembly level. These results highlight distinct performance characteristics across different categories of EMA methods. Consensus methods, which leverage inter-model agreement, typically achieve higher overall correlation across large model pools, whereas single-model methods function independently of ensemble-derived information and confer enhanced flexibility for real-world applications^19,65^. Notably, DeepUMQA-Global enables reliable top-model selection across multiple evaluation metrics using only a single input model, without reliance on cross-model consensus. This model-pool-independent evaluation strategy achieves competitive performance while maintaining high scalability, supporting its applicability for large-scale structural model accuracy estimation and downstream analyses.

### Performance Comparison of DeepUMQA-Global-Con with CASP16 EMA Methods

To further investigate the effectiveness and scalability of single-model strategies, we introduce DeepUMQA-Global-Con, a lightweight consensus-enhanced extension of DeepUMQA-Global. In this framework, DeepUMQA-Global is first used to rank candidate models, and the top five models are selected as a high-confidence reference subset. Structural alignments between each candidate model and the reference subset are then performed using US-align^66^, and the final global accuracy score is computed as the average structural similarity across these alignments. In contrast to conventional consensus approaches that often rely on extensive pairwise comparisons across the model pool^19,65^, this strategy requires only a limited number of alignments. We further benchmarked DeepUMQA-Global-Con against all the EMA methods reported in CASP16 on the CASP16 benchmark test set.

DeepUMQA-Global-Con achieved the best performance for the SCORE evaluation, attaining the highest overall Z-score of 167.102 and exceeding that of the second-ranked method by 39.845 units (**Fig. 4a**). It also achieved the highest Z-scores for the Pearson, Spearman, ROC AUC and Loss with respect to both the TM-score and Oligo-GDTTS, indicating improved robustness in global model ranking across diverse protein complexes (**Supplementary Table 15**). Consistent with this trend, DeepUMQA-Global-Con correctly identified the top-ranked model for 19 targets (Top1Loss<0.05), representing the highest performance among all the CASP16 EMA methods (**Fig. 4c**). These results suggest that incorporating a small, high-confidence reference subset enables more effective utilization of inter-model structural similarity without extensive pairwise comparisons across the full model pool. By anchoring consensus information to top candidates initially ranked by DeepUMQAÖGlobal, the approach reinforces consistent structural signals while reducing the influence of lower-quality or conformationally heterogeneous models. This strategy of combining single-model quality assessment and model consensus information likely contributes to the observed improvements in both overall ranking accuracy and the top-model selection.

**Fig. 4.**
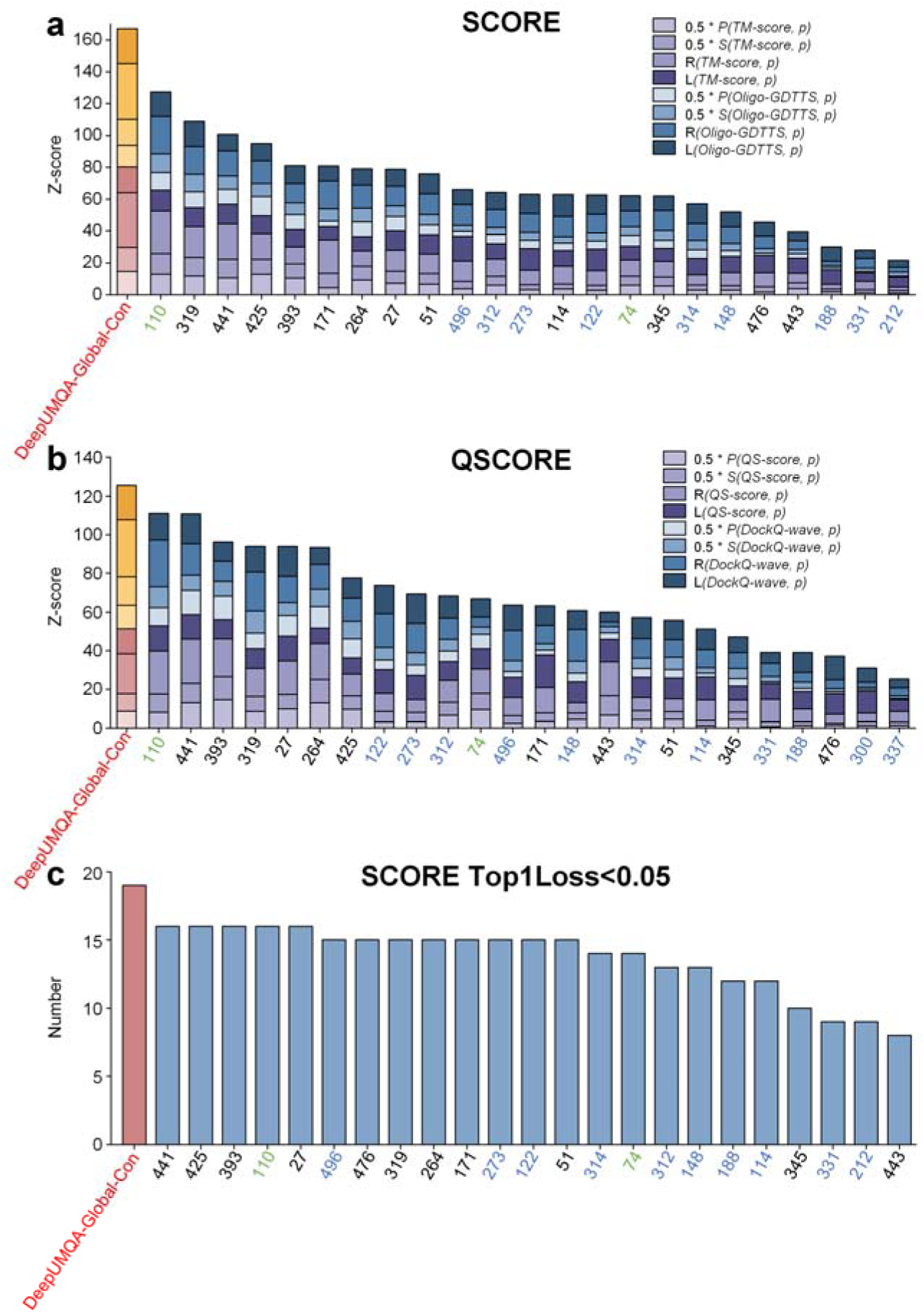
| Performance comparison of DeepUMQA-Global-Con with all CASP16 EMA methods. a, Performance comparison of DeepUMQA-Global-Con with all CASP16 EMA methods in the SCORE evaluation. **b,** Performance comparison of DeepUMQA-Global-Con with all CASP16 EMA methods in the QSCORE evaluation. **c,** Number of targets with a Top1Loss of <0.05 in the SCORE evaluation. All CASP16 EMA methods are distinguished by color, with consensus methods shown in black, quasi-single-model methods in green, and single-model methods in blue.

DeepUMQA-Global-Con also achieved the best performance in the QSCORE evaluation, attaining the highest overall Z-score among all methods and outperforming the second-ranked method by 25.997 units (**Fig. 4b**; **Supplementary Table 16**). Performance gains were more pronounced for the DockQ-wave than for the QS-score. DeepUMQA-Global-Con achieved the highest Spearman correlation, ROC AUC and Loss values with respect to DockQ-wave, whereas its performance improvements were relatively modest for the QS-score^17^. This difference is likely due to the distinct characteristics of the two interface reference metrics and their differential sensitivity to the consensus information derived from structural similarity to the high-confidence reference subset. DockQ-wave^17^ is a continuous variant of the DockQ^64^ metric for evaluating interface accuracy, which integrates the fraction of native contacts (F_nat_), interface RMSD (iRMSD) and ligand RMSD (LRMSD) into a unified score that captures both interfacial contact recovery and geometric deviations. As the proposed approach computes structural similarity between each candidate model and a high-confidence reference subset, it preferentially captures consistency in interface geometry and relative chain orientation, thereby contributing to the observed performance gains^17,64^. In contrast, the QS-score evaluates the agreement of interfacial residue–residue contacts and their topological organization, with a primary focus on contact-level correspondence rather than geometric alignment^17^. Consequently, improvements derived from structural alignment are less directly reflected in QS-score, resulting in relatively modest gains.

DeepUMQA-Global-Con consistently improves performance across ranking metrics, yielding enhanced stability, discriminative power and top-model selection accuracy. These gains can be attributed to DeepUMQA-Global, whose strong and reliable ranking capability enables the construction of a compact, target-specific reference subset. By capturing a set of structurally plausible conformations, this subset allows consensus information to emphasize consistent structural patterns while reducing the influence of lower-quality models. The aggregation of similarity scores over this subset reduces variance and improves the alignment between the predicted scores and reference accuracy. Importantly, this design confines structural comparisons to a limited set of high-confidence models, resulting in approximately linear computational scaling of O(N). In contrast, conventional consensus approaches typically rely on exhaustive pairwise comparisons across the full set of models, leading to quadratic scaling O(N²) and higher computational cost^65^. These results demonstrate that DeepUMQA-Global-Con, as a lightweight consensus-inspired approach, improves accuracy estimation while capturing consensus signals with reduced structural alignment. More importantly, they further validated the effectiveness and robustness of DeepUMQA-Global, and suggest that it can serve as a general and extensible module that may be readily integrated into other EMA methods.

### Ablation and Interpretability Analyses of DeepUMQA-Global

To systematically quantify the contributions of distinct features and network module components to global accuracy estimation in DeepUMQA-Global, we performed a comprehensive suite of ablation and interpretability analyses (**Fig. 5**). All analyses were implemented on the CASP16 benchmark test set, for targets with sequence lengths less than 3,000 residues. We first evaluated the performance changes upon ablation of individual features, enabling quantification of the relative importance of different features for global accuracy estimation. We subsequently performed ablation of the core network modules to identify their indispensable roles in capturing global structural patterns and ensuring robust estimation of global model accuracy. Correlation analyses were performed within the residue-wise and inter-residue feature representations to assess internal redundancy. Finally, we used Uniform Manifold Approximation and Projection (UMAP)^67^ to visualize the evolution of learned embeddings across network modules, examining how the model progressively encodes and aggregates heterogeneous inputs for global accuracy estimation. We first evaluated the impact of removing individual features on global fold and interface accuracy estimation in the SCORE and QSCORE evaluations (**Fig. 5a**). Among all features, SASP emerged as the dominant contributor. Its removal led to substantial performance degradation, with overall Z-scores decreasing from 101.340 to 60.420 (−40.920) in the SCORE evaluation and from 93.141 to 47.827 (−45.314) in the QSCORE evaluation, indicating that the SASP provides critical topological priors for accurate global model evaluation. SAGS also contributed substantially to performance, with its removal reducing Z-scores to 82.407 (−18.933) for SCORE and 75.858 (−17.283) for QSCORE, suggesting that SAGS encodes structure-aware sequence preferences that support fold-level accuracy estimation. The SASP (single-template) variant, which infers topological priors from only one structural template instead of multiple templates, with Z-scores decreasing to 86.118 (−15.222) for SCORE and 82.294 (−10.847) for QSCORE, highlighting the importance of aggregating multiple template-derived structural priors. By comparison, the removal of SALS, as well as the combined removal of SASA and VTIO, resulted in moderate performance reductions, indicating that these features play auxiliary roles in supporting global accuracy estimation. Detailed results are provided in **Supplementary Tables 17-18**.

**Figure 5.**
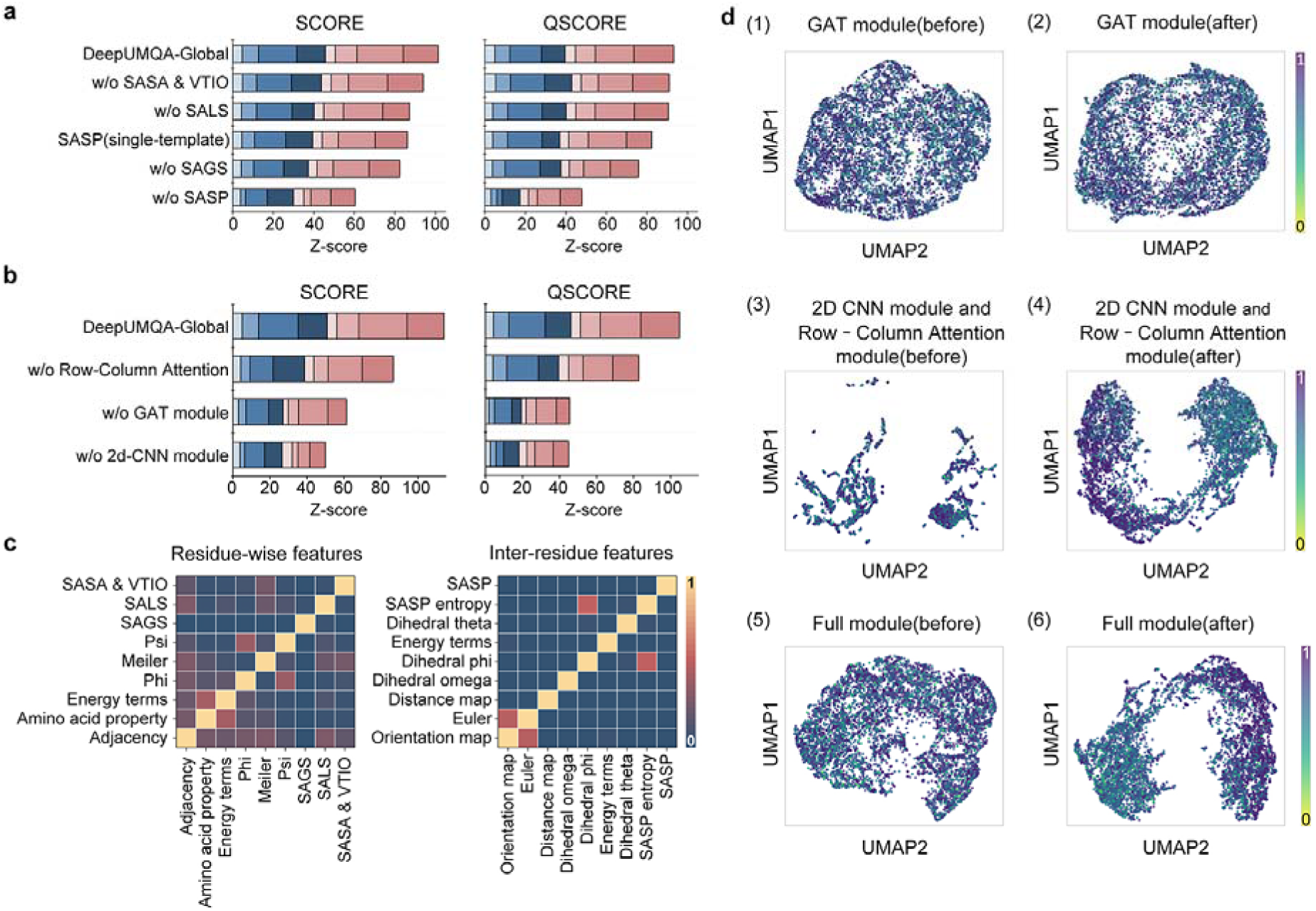
A**b**lation **and interpretability analyses of DeepUMQA-Global. a,** Feature ablation results showing the performance of removing individual features in the SCORE (left) and QSCORE (right) evaluation. **b,** Network ablation results evaluating the performance of removing the Row–Column Attention module, the GAT module, and the 2D CNN module. **c,** Feature correlation matrices for residue-wise features (left) and inter-residue features (right), illustrating inter-feature linear correlations. **d,** UMAP projections visualizing the evolution of learned representations across successive network stages: before and after the GAT module (1,2); 2D CNN module and Row–Column Attention module (3,4); as well as GAT, 2D CNN and Row–Column Attention module (5,6).

We next evaluated the contributions of core network modules by ablating the GAT module, 2D CNN module, or Row–Column attention module (**Fig 5b**). Among these architectural components, ablation of the 2D CNN module resulted in the most pronounced performance degradation, with overall Z-scores decreasing from 114.000 to 49.922 (−64.078) in the SCORE evaluation and from 104.825 to 44.902 (−59.923) in the QSCORE evaluation. This finding indicates that the 2D CNN^57^ module is indispensable for integrating pairwise structural consistency and capturing global topological patterns, which are essential for accurate global model accuracy estimation. Ablation of the GAT^56^ module also resulted in substantial performance degradation, with Z-scores decreasing to 61.414 (−52.586) for SCORE and 82.839 (−21.986) for QSCORE, demonstrating its critical role in aggregating residue-wise context information and propagating local structural dependencies for reliable fold and interface assessment. Similarly, removing the Row–Column attention^58^ module caused notable declines in predictive performance, with Z-scores decreasing to 86.701 (−27.299) for SCORE evaluation and 45.410 (−59.415) for QSCORE evaluation, highlighting its importance in adaptively modeling long-range inter-residue dependencies. Detailed results provided in **Supplementary Tables 19**–**20**. These results demonstrate that each module plays a distinctive, non-redundant role in DeepUMQA-Global. The consistent performance deterioration across the SCORE and QSCORE evaluations underscores the necessity of synergistically integrating these complementary module designs for robust global fold and interface accuracy estimation.

Correlation analysis further supported the ability of DeepUMQA-Global to integrate non-redundant features (**Fig. 5c**). At the residue-wise level, SAGS, SALS, and physicochemical features including SASA and VTIO all exhibit pairwise linear correlations no higher than 0.30, with SAGS in particular showing near-zero correlation (|r|≤0.05) with all other residue-level inputs, indicating that the residue-level features capture largely independent information. At the inter-residue level, most features exhibit weak intercorrelations, with two notable exceptions: the SASP entropy features show a moderate correlation with the φ backbone dihedral angle (r = 0.49), and the Euler angles are moderately correlated with inter-residue orientation maps (r = 0.45). All other pairwise feature correlations are less than 0.04, reflecting that the integrated pairwise feature set captures distinct, largely non-redundant facets of residue–residue interactions. These results demonstrate that DeepUMQA-Global effectively integrates complementary, weakly dependent features, supporting robust global model accuracy estimation.

To investigate how DeepUMQA-Global progressively organizes structural information to discriminate model quality, we visualized the evolution of learned representations via UMAP^67^ across successive network stages (**Fig. 5d**). After the GAT module processed the of residue-level features, the embeddings largely overlapped, with no clear separation by target identity or model quality. This finding indicates that, at this stage, residue-level representations primarily encode local structural context but lack discriminative signals for model quality. Following the application of the 2D CNN and Row–Column attention modules to pairwise representations, the latent space became strongly structured by model accuracy, with high-quality models from different targets clustered together, whereas low-quality models segregated into distinct regions. Accuracy-aligned clustering was further consolidated, with clear separation between high- and low-quality models persisting in UMAP visualization. Collectively, these UMAP visualizations analyses demonstrate that DeepUMQA-Global enables progressive refinement of its learned representations through the synergistic integration of residue-level and pairwise structural information. These modules work together to enhance signals related to model accuracy while reducing protein-specific differences, producing robust quality representations that generalize across different protein targets and folds, highlighting the rational hierarchical design of the network framework for robust and generalizable global model accuracy estimation.

Collectively, these analyses demonstrate that DeepUMQA-Global integrates complementary signals from sequence, physicochemical, and structural features, each contributing non-redundant information relevant to global model accuracy. Network ablations delineate the functional roles of major modules, whereas correlation and embedding analyses reveal weak inter-feature correlations and a progressive structuring of latent representations by model quality. These results indicate that DeepUMQA-Global hierarchically organizes multi-scale cues to form internal representations that robustly reflect fold- and interface-level accuracy without over-relying on any single feature source.

### Exploration of DeepUMQA-Global for Model Accuracy Estimation on Alternative Conformation States

Proteins adopt multiple alternative conformation states that are critical for their biological function^68^. However, most existing EMA methods focus on static protein structures and do not explicitly account for relationships among these distinct states^69^. Consensus methods assess model accuracy by aggregating pairwise structural similarities across a model pool, under the assumption that higher-quality models tend to show greater structural consistency. For proteins with multiple conformations, this strategy may become less effective, as structurally divergent yet functionally relevant conformations may fail to form coherent clusters. This can weaken consensus signals and introduce biased quality estimates. Single-model methods derive quality estimates from individual model representations. Their predictions often become unstable for proteins that populate alternative conformational states due to the lack of contextual information across structural ensembles^6,7,17,29^. These issues highlight that accurate evaluation of such alternative conformational states remains an open problem. To explore whether DeepUMQA-Global can reliably evaluate model accuracy for each conformational state and discriminate high-quality models from a mixed pool containing two states, we selected target T1249 for testing, the only alternative conformational protein complex target in the CASP16 EMA benchmark ^69^. The other alternative conformational states target T1294 was excluded due to its two structural states exhibit only subtle differences with negligible domain rearrangement, which were deemed insufficient to represent meaningful alternative conformations^69^. To further systematically validate the generality and consistency of DeepUMQA-Global for evaluating model accuracy across alternative conformations, we evaluated it on 91 apo–holo pairs from the CoDNaS^70^ database, which encompass structurally divergent yet functionally meaningful native states of the same protein.

Target T1249 is a trimeric spike protein of the Sabiá virus, an arenavirus associated with Brazilian hemorrhagic fever^69^. Cryo-EM structures have resolved two alternative conformational states: T1249v1 represents a compact, tightly packed closed state, while T1249v2 adopts an open state induced by acidic pH and stabilized by metal ion coordination^69^ (**Fig. 6a (1)**). Interconversion between these states involves concerted rigid-body rearrangements that alter protomer packing and inter-subunit interfaces^69^. We observed that when all submitted T1249v1 and T1249v2 models were evaluated against both experimental reference structures, the TM-score and DockQ-wave formed continuous, overlapping distributions with no clear bimodal separation between the two conformational states (**Fig. 6a (2)**). Consistent with the CASP16 assessment, the models generally achieved higher accuracy for the closed T1249v1 state than for the open T1249v2 state, reflecting a marginal global bias toward the closed conformation^69^. The overlap between alternative conformational states complicates the discrimination of high-quality models across distinct states, highlighting the critical need for accurate model accuracy estimation. The red-highlighted points in **Fig. 6a (2)** denote the top-ranked models selected for our subsequent evaluation of DeepUMQA-Global.

**Figure 6.**
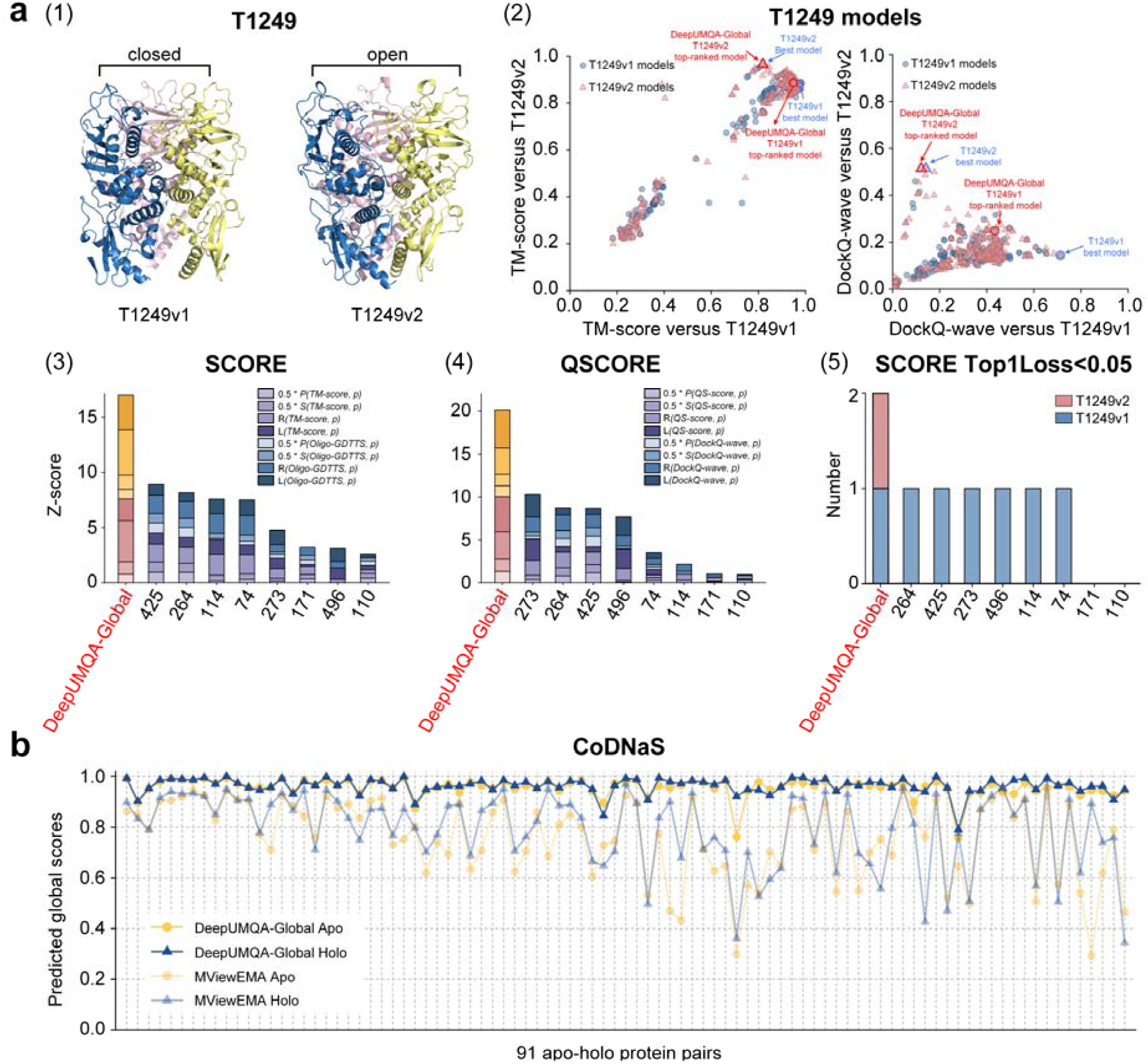
Performance of DeepUMQA-Global for model accuracy estimation on alternative conformational states. a, (1) The CASP16 target T1249 was released in two alternative conformational states, with T1249v1 representing the closed state and T1249v2 representing the open state. (2) TM-score and DockQ-wave distributions of all the T1249v1 and T1249v2 models. Each model was aligned versus both the T1249v1 and T1249v2 reference structures, generating two TM-score and DockQ-wave distributions that distinguished models adopting the closed state (blue) from those adopting the open state (red). The models highlighted by red circles and red triangles correspond to the top-ranked T1249v1 and T1249v2 models selected by DeepUMQA-Global, while the blue highlight denotes the best model for each conformational state. (3) Performance comparison of DeepUMQA-Global with all EMA methods reported in CASP16 on T1249v1 and T1249v2 for SCORE. (4) Performance comparison of DeepUMQA-Global with all EMA methods reported in CASP16 on T1249v1 and T1249v2 for QSCORE. (5) Number of T1249v1 and T1249v2 targets for SCORE with Top1Loss<0.05. **b.** Performance comparison of DeepUMQA-Global with MVEMA for 91 CoDNaS apo–holo protein pairs. Dark yellow circles denote global quality scores predicted by DeepUMQA-Global for apo structures, and dark blue triangles denote corresponding scores for holo structures. Light yellow circles and light blue triangles represent global quality scores predicted by MViewEMA for apo and holo structures, respectively.

We compared the performance of DeepUMQA-Global with CASP16 EMA methods on the alternative conformation target T1249 (**Fig. 6a**, **Supplementary Fig. 3**). In the SCORE evaluation (**Fig. 6a (3)**), DeepUMQA-Global achieved the highest overall Z-score for T1249v1 and T1249v2 (16.983), outperforming the second-ranked method by 8.066 units (**Supplementary Table 21**). On the open state T1249v2, it achieved the highest Z-score of 11.566, which exceeded that of the second-ranked method by 7.997 units (**Supplementary Table 22**). On the closed state T1249v1, it remained competitive, ranking third among all methods (**Supplementary Table 23**). Notably, DeepUMQA-Global was the only method with a SCORE Top1Loss < 0.05 for both alternative conformation states, highlighting its potential for top-ranked model selection of alternative conformational states (**Fig. 6a (5)**). To further test its performance under mixed-state conditions, we pooled models from both conformations and assessed whether DeepUMQA-Global could correctly prioritize the most accurate models as measured by the TM-score. Among its top two ranked models, one corresponded to T1249v1 and the other to T1249v2. Importantly, the top-ranked model for T1249v2 matched the ground-truth best model of that state, and the selected T1249v1 model showed only a marginal deviation from the optimal reference (Top1Loss = 0.016). In the QSCORE evaluation (**Fig. 6a (4)**), we observed a consistent trend. DeepUMQA-Global achieved the highest overall Z-score for T1249v1 and T1249v2 (20.123), outperforming the second-ranked method by 9.824 units. On the open state (T1249v2), the Z-score was again the highest (12.851), outperforming the other methods (4.727 units) and ranking second on T1249v1 (**Supplementary Tables 24–26**). These results demonstrate that DeepUMQA-Global can not only accurately evaluate model accuracy for individual conformational states, but also shows potential for discriminating near-native models across both alternative conformational states.

To further validate the generality of DeepUMQAÖGlobal and its ability to provide stable, reliable accuracy estimates for alternative conformational states, we systematically assessed 91 apo–holo protein pairs from the CoDNaS database (**Fig. 6b**). The apo–holo pairs in **Fig. 6b** are arranged from left to right following the column-wise order of **Supplementary Table 27**. As shown in **Supplementary Fig 4**, DeepUMQA-Global yielded highly consistent scores across the two alternative states of each protein pair, with a small apo–holo difference (mean = 0.013; 95% confidence interval: 0.009–0.017). In contrast, MViewEMA^71^, a single-model EMA method we previously developed and participated in CASP16 (group 148), exhibited larger apo–holo discrepancies (mean = 0.0690; 95% confidence interval: 0.052–0.090), indicating sensitivity to conformational variation. The difference in apo–holo score deviations between the two methods was statistically significant as assessed by a two-sided Wilcoxon signed-rank test (*P* = 2.679× 10⁻¹²). These observations indicate that DeepUMQA-Global may yield consistent and reliable accuracy estimates across both conformational states, without a pronounced preference for either native state.

The potential of DeepUMQA-Global to accurately estimate alternative conformational states may be derived from the design of a structure-sequence crossÖvalidation mechanism. SASP encodes a spatially probability distribution over residue–residue relative positioning, capturing conserved global fold topology while tolerating rigid-body rearrangements and subtle conformational shifts, which may enhance its reliability for evaluating conformationally dynamic proteins. SAGS captures global sequence preferences from three-dimensional structure, encoding residue-level sequence fitness that remains stable evolutionary constraints across alternative conformational states. Their integration may enable DeepUMQA-Global to provide stable accuracy estimates with favorable consistency across distinct alternative conformations, supporting its application in the study of alternative conformation states systems.

## Conclusions

Accurate estimation of protein model accuracy is a fundamental step in computational structural biology, as it provides critical guidance for model selection, functional inference, and rational design^6,7,17,21,29^. In this study, we present DeepUMQA-Global, a single-model EMA method that exploits sequence–structure bidirectional compatibility. By implicitly encoding consistency between structure-aware sequence and the amino acid sequence, as well as consistency between sequence-aware structural profile and the three-dimensional structure, DeepUMQA-Global enables accurate evaluation of model accuracy.

The results demonstrate that DeepUMQA-Global provides consistent and reliable accuracy estimates for protein structural models. Notably, DeepUMQA-Global outperforms AlphaFold3^9^ self-assessment in the Pearson and Spearman correlation coefficients, indicating that DeepUMQA-Global captures structural accuracy signals beyond those encoded in self-assessment methods, and may serve as a valuable external complement for confidence scores of predictors. In retrospective evaluations on the CASP16^31^ benchmark test set, DeepUMQA-Global surpasses CASP16 single-model methods and achieves a performance comparable to consensus-based methods, demonstrating its ability to reliably evaluate model accuracy. Furthermore, the performance of our lightweight consensus strategy underscores the effectiveness and scalability of DeepUMQA-Global, highlighting its potential as a general, extensible building block for integration into other EMA pipelines. Interestingly, DeepUMQA-Global demonstrates the potential in providing stable accuracy estimates for alternative conformational states, as well as may reliable discriminate between two alternative conformations within heterogeneous structural ensembles.

Looking ahead, several directions may further improve the applicability of DeepUMQA-Global. Future extensions could incorporate dynamic and functional attributes, such as binding affinities and interaction propensities, to enable a more holistic evaluation of dynamic protein systems^72^. Expansion of the framework to multicomponent biomolecular assemblies, including protein–DNA, protein–RNA, and ribonucleoprotein complexes would broaden its applicability beyond protein–protein complex^9,73^. Further integration with experimental data from cryo-EM density maps^74^ and X-ray crystallographic^75^ may enable reciprocal refinement between computational predictions and experimental observations, leading to more reliable structural interpretations. Collectively, the results of this work establish DeepUMQA-Global as an accurate, generalizable, and extensible single-model EMA approach, with broad utility for reliable accuracy evaluation in protein structural modeling.

## Methods

This section provides a detailed introduction to the technical details of DeepUMQA-Global, specifically covering the construction of training and test datasets, the extraction methods for three complementary protein model features, the deep learning network architecture, and the complete model training process.

### Dataset

#### Train set

The training dataset for DeepUMQA-Global was constructed from the Protein Data Bank (PDB)^60^, following the training set construction strategy of DeepAccNet^45^. First, proteins with experimentally resolved structures at a resolution of 2.5 Å and a length of 50–400 residues were selected from the PISCES server (data version March 2022). Then, redundancy was removed using MMseqs2^76^ with a 40% sequence identity cutoff and 80% reciprocal sequence coverage, yielding a set of 7,992 non-redundant monomeric protein targets. To ensure no overlap with the test datasets, complexes from both CASP16 and the docking benchmark were first split into individual chains, which were then compared against the training dataset. Any training targets showing similarity above the same redundancy thresholds were excluded using MMseqs2^76^, resulting in a final non-redundant training set comprising 7,840 monomeric protein targets, among which 3,583 were single-domain proteins and 4,257 were multi-domain proteins, as classified using Merizo^77^ domain segmentation. To generate diverse training examples with a wide range of structural accuracy, we produced multiple structural decoys for each target using comparative modeling, native structural perturbation, and deep learning-based folding methods. On average, ∼140 decoys were generated per protein, resulting in a total of 1,065,994 structural decoys. The entire dataset was randomly split at a 9:1 ratio, with 1,024,863 decoys allocated to model training and 41,131 to validation.

#### Test sets

The test sets include three protein complex datasets: the AlphaFold3 test set, the docking test sets, and the CASP16 benchmark test set. These datasets were collected after January 2024 and are completely separated in time from the training set (which includes data prior to March 2022), ensuring unbiased and reliable performance evaluation. For the AlphaFold3 test set, all targets were released after the AlphaFold3 training data cutoff date of 30 September 2021, which prevents potential evaluation bias due to data leakage from AF3’s training set.

##### AlphaFold3 test set

The AlphaFold3 test set was constructed by selecting 26 protein complex targets from CASP16, each with a sequence length not exceeding 3,000 residues. For each target, 50 models were generated using the AlphaFold3 online server (https://alphafoldserver.com/) under stochastic sampling with random seeds, resulting in a total of 1,300 models across all targets. Two CASP16 targets (T1249 and T1294), which exhibit experimentally observed conformational heterogeneity, were excluded to reduce potential ambiguity arising from conformational variability, given that AlphaFold3 predictions typically represent a single dominant conformational state. The full list of AlphaFold3 test set targets is provided in **Supplementary Table 28**.

##### Docking-based test set

We selected protein complex structures released in PDB^60^ between January 1, 2024, and April 1, 2024. These complexes were required to have a resolution better than 2.5 Å and chain lengths ranging from 20 to 1000 residues. Sequence redundancy was removed at 40% identity using MMseqs2^76^, ensuring non-redundancy both within the dataset and relative to the training set. After filtering, 107 complexes were retained as benchmark targets. For each complex, HDOCK and DiffDock-PP were independently applied to perform docking re-modeling, generating 200 candidate models per target. The full list of this test set targets is provided in **Supplementary Table 29**.

##### CASP16 benchmark test set

The CASP16 benchmark test set was derived from the model sets released by the CASP16 organizers for EMA evaluation. The dataset comprises models submitted by participating structure prediction groups for protein complex targets under the CASP16 EMA QMODE1 framework. A total of 38 targets were included for SCORE evaluation and 39 targets for QSCORE evaluation^16^, with approximately 350 models per target. Two targets (H1229 and H1230) were excluded due to the absence of experimentally resolved reference structures in the official assessment^16^. The targets whose sequence lengths less than 3,000 residues were retained due to computational constraints. This filtering resulted in a test set of 30 targets, comprising a total of 10,521 models for evaluation. The complete list of targets is provided in **Supplementary Table 30**.

### Input Features

For each input model and sequence, DeepUMQA-Global extracts a comprehensive set of features from three complementary perspectives, including structural, sequence, and physicochemical features. By enabling implicit bidirectional inference between sequence and structure, these features capture multiscale consistency to support reliable and robust global model accuracy estimation. Detailed shapes and descriptions are provided in **Supplementary Table 1**.

1. Structural At the structure level, we extract template-derived sequence-aware structural profiles (SASP) and inter-residue features to assess the plausibility of global fold topology. For SASP, a corresponding high-quality reference structure was first generated from its amino acid sequence using AlphaFold3 and used as the query for structural template search. Subsequently, Foldseek-Multimer^52^ was employed to search the official PDB100 database (PDB snapshot downloaded in November 2023), template complexes with high global conformational similarity to the query structure were obtained. To prevent information leakage during training, any templates with 100% sequence identity to the query were strictly excluded. When more than 300 templates were retrieved, only the 300 top-ranked templates were retained for subsequent analysis. The query structure was then structurally aligned to each selected template using US-align^66^. For the aligned regions, residue-residue geometric relationships were extracted by computing the distances between *C_β_* atoms (*C_α_* for glycine) for every residue pair (*i*, *j*) in each aligned template. To ensure a consistent representation across templates, distances were restricted to the range of 2–20 Å and discretized into 36 uniform bins with a bin width of 0.5 Å. For each residue pair (*i*, *j*), the distances observed across all aligned templates were accumulated into a histogram over the predefined bins and normalized to obtain a discrete distance distribution. This distribution summarizes how frequently a given residue–residue separation is supported by the template set and constitutes the structural profile feature for that residue pair. To further quantify the variability of residue–residue geometries across templates, an entropy term was computed for each residue pair based on its distance distribution 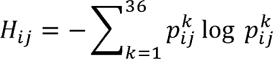, where 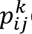 denotes the normalized frequency of distances falling into the *k*-th bin. Higher entropy values indicate more dispersed distance distributions and weaker structural constraints, whereas lower entropy reflects stronger geometric conservation across templates. By jointly incorporating the residue–residue distance distributions and their associated entropy, the structural profile encodes template-derived geometric priors that capture conserved and variable topological relationships beyond a single structure, providing a quantitative and interpretable representation of global fold plausibility. In addition, we incorporate conventional pairwise descriptors from the input structural model, including inter-residue distance and orientation maps.
2. Sequence At the sequence level, we extract features including structure-aware global sequence (SAGS), structure-aware local sequence (SALS), and amino acid sequence of the input model to evaluate the consistency between the input sequence and the sequence preferences implicitly encoded by its structural environment. SAGS is derived from the intermediate layer embeddings of ProteinMPNN^50^. ProteinMPNN takes the three-dimensional structure of a protein model as conditional input, constructs local geometric embeddings using backbone atom coordinates, and encodes and aggregates atomic-level spatial relationships through a graph neural network to form residue-centered structural environment embedding that describes how a fixed conformation restricts the amino acid choices that can be accommodated within each local structural context^50^, thereby defining a feasible set of sequences compatible with the overall fold. We use this embedding to quantify the extent to which the input sequence departs from these constraints, thus providing a global signal of sequence–structure consistency. SALS feature is derived from 3Di structural sequence embedding generated by Foldseek^51^. Based on the local geometric configuration of the protein backbone and spatial orientation of neighboring residues, Foldseek maps continuous three-dimensional structural information into a set of discrete 3Di structural states, thereby encoding the local conformational type of each residue in a sequential format. Each 3Di state corresponds to a class of canonical backbone geometric configurations and relative orientation patterns recurrent in known protein structures, which serve to delineate the conformational category of individual residues within their local structural microenvironment^51^. In this study, 3Di structural sequences are converted into one-hot encoded vectors for use as input features, enabling characterization of the conformational category distribution of predicted models at the local scale. As the 3Di alphabet is derived from statistical learning on large repositories of known protein structures, this embedding reflects the relationship between the locally predicted geometric conformations and conformational categories prevalent in natural protein structures. We incorporate this 3Di structural sequence feature to enable the model to discern the conformational pattern composition of predicted structures at the local scale, thereby providing complementary information for assessing the plausibility of local structural conformations.
3. Physicochemical At the physicochemical level, we extracted SASA and VTIO descriptors together with a set of physicochemical properties to characterize residue exposure, local contact geometry, and intrinsic energetic context. The SASA descriptor quantifies the surface area of each residue derived from Voronoi tessellation, including contributions from contacts with neighboring residues as well as exposure to the surrounding solvent^46,53^. VTIO encodes the local directional arrangement of residue contacts by summing the normal vectors defined between the atoms of each residue and the atoms of neighboring residues. Additionally, we extracted general physicochemical properties, including meiler^45^, amino acid properties, BLOSUM62^54^, Rosetta one-body and two-body energy terms^55^, backbone bond lengths and angles, torsion angles, Euler angles and voxelization^45^.

### Network Architecture

DeepUMQA-Global employs a hybrid deep-learning architecture that integrates residue-level and residue–residue pairwise representations to estimate global protein structure quality. Given an input structure, residue-wise and inter-residue features are first constructed to encode sequence-derived and structure-derived information. The residue-wise features are processed by a graph attention network (GAT) module to generate node embeddings, while the inter-residue features are encoded using a two-dimensional convolutional neural network (2D CNN) module together with a Row–Column attention module to capture geometric relationships across residue pairs. The node and pair representations are then integrated through a gating mechanism that couples single-residue and pairwise information. The fused representations are aggregated to produce global quality estimates, including a global score (pScore) for the full structure and an interface-specific score (ipScore) computed by restricting residues to protein–protein interface regions. The detailed design of each module is described in the following sections.

1. GAT Module In the GAT module, residue-wise features are encoded as node attributes in a graph representation. These node features are constructed by combining sequence-derived embeddings with residue-level physicochemical descriptors, including amino acid properties and voronoi tessellation-derived geometric features. Edges are defined between residue pairs whose *C_α_* atoms are within a 15 Å cutoff based on pairwise *C_α_*–*C_α_* distances. The node and edge definitions form a residue-wise 1D embeddings. The 1D embedding is first processed by a self-attention graph pooling (SAGPooling)^78^ layer with a sampling ratio of 0.5 to retain the most informative residues while reducing the graph size, and then refined through four consecutive graph attention (GAT)^56^ layers (eight attention heads, dropout = 0.25, GELU^79^ activation). A final linear projection maps the aggregated embedding to a single representation.
2. 2D CNN and Row–Column Attention Module Structural features and inter-residue physicochemical features are encoded as two-dimensional (2D) embeddings. Then, 2D embeddings are processed through the 2D CNN^57^ module, which applies successive convolutions with normalization and Gaussian error linear units (GELU)^79^ activation to extract local structural patterns and compress channel dimensions while enhancing representational compactness. Spatial resolution is reduced by a downsampling block that applies a 3×3 convolution neural network with stride of 2, aggregating contextual information while compressing spatial dimensions and retaining salient structural signals. The downsampling embeddings are then refined using a Row–Column attention^58^ module, implemented as a single-layer Pair2Pair block^80^ with four attention heads, 128-dimensional feature channels, a feedforward expansion ratio of 2, and 0.1 dropout, capturing long-range correlations and evaluating directional dependencies along both residue indices. Finally, a 1 × 1 convolution produces the pair representation.

The single representation is broadcast and applied via element-wise multiplication to gate the pairwise representation, and the resulting gated representations are averaged over both rows and columns to generate the global accuracy score (pScore) and the interface-specific global score (ipScore).

### Training

To objectively quantify global structural similarity, we used the TM-score, a normalized metric that evaluates the overall topological consistency between predicted and reference structures and is sensitive to global fold organization and relative chain positioning, to train DeepUMQA-Global on protein monomer datasets for predicting global model accuracy; the precise definition of the TM-score is provided in **Supplementary Note 1**. Selecting monomeric structures enables the preservation of multi-domain fold organization, while the inter-domain spatial relationships within monomers provide an implicit reference for the general geometric and physicochemical constraints governing chain–chain interactions, thereby supporting the learning of generalizable structural interaction principles in protein complexes. The network was implemented in PyTorch with PyG, and optimized using the Adam algorithm with an initial learning rate of 1×10⁻O, which decayed by a factor of 0.99 at each step. Model optimization employed the log-cosh loss:

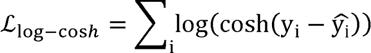

where *y_i_* denotes the true TM-score, *̂y_i_*_i_ is the network prediction, and *cosh* is the hyperbolic cosine function. This loss function behaves like the mean squared error for small residuals while approximating the mean absolute error for larger deviations, thereby combining smooth gradient propagation with robustness to outliers. Data were accessed via a customized dataset loader that performed random sampling of decoy structures within each training batch to ensure exposure to diverse residue compositions and structural contexts. Training and validation were performed using PyTorch Lightning, and the model with the lowest validation loss was selected as the checkpoint for downstream evaluation. All the experiments were conducted on a single NVIDIA A100 GPU, and training required approximately two days.

## Availability of data and materials

The authors declare that the data supporting the results and conclusions of this study are available within the paper and its Supplementary Information. Protein structure data were obtained from the Protein Data Bank (PDB) (https://www.rcsb.org). CASP16 EMA data, including predicted models, experimental structures, and assessment results, are available at https://predictioncenter.org/casp16/index.cgi. Structural models for the AlphaFold3 test set were generated using the online AlphaFold3 server (https://alphafoldserver.com/). Native protein structures from the CoDNaS dataset are available at http://ufq.unq.edu.ar/codnas/. CASP16 evaluations were performed using the publicly available CASP16 EMA evaluation pipeline (https://git.scicore.unibas.ch/schwede/casp16_ema). The PDB100 database is available at (https://steineggerlab.s3.amazonaws.com/foldseek/pdb100.tar.gz). The dataset for this study is publicly available via GitHub (https://github.com/iobio-zjut/DeepUMQA-Global/).

## Code availability

DeepUMQA-Global is available for download via GitHub at https://github.com/iobio-zjut/DeepUMQA-Global/. The online server of DeepUMQA-Global is made freely available at http://zhanglab-bioinf.com/DeepUMQAGlobal/.

## Consent for publication

Not applicable.

## Contributions

G.Z. conceived and supervised the research. G.Z., L.X., E.Y., and H.W. designed the experiments. L.X., E.Y.,H.W., T.Z., Q.Z., F.L., and D.L. collected the data and performed the experiments. G.Z., L.X., E.Y., and H.W. analyzed the data. L.X., E.Y., H.W., and G.Z. wrote the manuscript, and all the authors read and approved the final manuscript.

## Ethics declarations

### Ethics approval and consent to participate

Not applicable.

